# No Cost of Complexity in Bacteriophages Adapting to a Complex Environment

**DOI:** 10.1101/434324

**Authors:** Andrew M. Sackman, Darin R. Rokyta

## Abstract

A longstanding prediction in evolutionary biology is that organisms experience a so-called “cost of complexity” manifested as a decreasing rate of adaptation in populations as organisms or selective environments become increasingly complex. This theory assumes the ubiquity of antagonistic pleiotropy, or tradeoffs in fitness, for mutations affecting multiple traits or phenotypes. A particular manifestation of antagonism thought to be at play in adaptive dynamics involves the relationship between viral growth rate and capsid stability, an interaction that may impede the adaptation of viral pathogens to novel hosts and environments. Here, we present a comparison of the genetics of adaptation for populations of bacteriophages undergoing complete adaptive walks under both simple and complex selective conditions, with complexity being determined by the number of traits under directional selection. We found no evidence for a long-term cost of complexity in viruses experiencing complex selection, with on average at least as great a rate of adaptation under more complex conditions, and rampant evidence for synergistic, rather than antagonistic, pleiotropy. The lack of evident tradeoffs between multiple phenotypes implies that emerging pathogens may be able to improve their growth in many different hosts or environments simultaneously and to do so at a faster rate than previously anticipated.

## Introduction

Adaptation is a process by which a population moves toward a multi-phenotypic optimum resulting in an increase in fitness, and the underlying genetic architecture of traits strongly determines evolutionary outcomes (Fisher 1930; Otto 2004; Wright 1969; Weinreich et al. 2005). Fisher’s geometric model characterizes adaptive evolution by the movement of a population through a phenotypic space towards a fitness optimum (Fisher 1930). The dimensionality of the phenotypic space is determined by the number of traits under selection. The model allows for universal pleiotropy, or, that any mutation can affect any number of traits with a beneficial or deleterious effect of any magnitude. A seemingly uncontroversial prediction of the model is that the rate of adaptation, or the average rate of fitness increase per unit of time, is inversely related to the number of traits under selection (Orr 2000; Welch and Waxman 2003), meaning that complex organisms with a large number of traits under selection suffer a cost of complexity in the form of a reduced rate of adaptation relative to simpler organisms. This prediction results from antagonistic pleiotropy, or tradeoffs between competing traits under selection.

Pleiotropy is characterized by a single gene or locus having an effect on multiple phenotypes or traits (Otto 2004; Wagner and Zhang 2011; Østman et al. 2011). Antagonistic pleiotropy occurs when a mutation that is beneficial for one trait incurs a cost for one or more additional traits (Mather and Harrison 1949; Cooper et al. 2001; Bono et al. 2017), while synergistic pleiotropy occurs when a single mutation simultaneously improves two or more traits (McGee et al. 2016). Synergistic pleiotropy, therefore, could accelerate the rate of adaptation for complex organisms, as mutations would benefit multiple traits simultaneously. Antagonistic pleiotropy has been implicated in phenomena such as the evolution of senescence (Williams 1957; Rose and Charlesworth 1980), the emergence of cooperation (Foster et al. 2004), and niche evolution (Kassen 2002; MacLean et al. 2004), but the purported cost of complexity has not often been empirically tested owing to the difficulty of calculating rates of adaptation over long evolutionary timescales in organisms or systems of measurable complexity.

A particular constraint that may be imposed by antagonistic pleiotropy involves the evolution of viral growth rate and capsid stability (McGee et al. 2016). Most mutations improving protein function or imbuing new functions are destabilizing (DePristo et al. 2005; Tokuriki et al. 2008; Tokuriki and Tawfik 2009; Phillips et al. 2017; Geller et al. 2018), especially when multiple functional mutations fix in succession (Bloom et al. 2005; gong et al. 2013). Furthermore, the ideal range of contact energy between capsid subunits for proper assembly may be narrow, as weak contact energy prevents assembly and strong contact energy can promote kinetic traps, resulting in many partially formed capsids and few free subunits available to complete assembly (Zlotnick 1994; Ceres and Zlotnick 2002; Zlotnick 2003, 2005). The geometry of the viral capsid amplifies the potential of single mutations to improve the stability of a capsid and alter the binding energies of the capsid subunits, allowing individual mutations to have dramatic impacts on capsid assembly and stability.

McGee et al. (2016) found that single mutations could improve both growth rate and capsid stability in microvirid phage ID8 when selection acted strongly on both traits. However, tradeoffs emerged when selection acted only one trait or only weakly on the secondary trait, and the average growth-rate effects of individual synergistically pleiotropic mutations were significantly smaller than for mutations that fixed in a simple environment selecting only on growth rate. This previous work focused only on first-step beneficial mutations for a single genotype, and it remains unknown what constraints pleiotropy between growth rate and capsid stability may impose on the ability of viruses to simultaneously optimize both traits over a complete bout of adaptation to a novel environment (Goldhill and Turner 2014). Viruses encounter a wide variety of potential hosts and environmental conditions before, during, and after infection, and the degree to which viruses incur a cost of complexity will greatly impact the rate at which they may evolve infectivity or virulence in new hosts.

We performed experimental adaptive walks (the sequential fixation of novel beneficial mutations in a population moving toward an optimum from a point some distance from the optimum) for two replicate lineages each of two pairs of unadapted, wild-type bacteriophages and corresponding growth-adapted versions of the same strains. The growth-adapted strains had been previously optimized under simple growth rate selection by Rokyta et al. (2009), exhausting the supply of accessible singlestep genetic variation benefiting growth rate. Complex selection was induced on each lineage by alternating periods of serial flask passaging where phage experienced selection on growth rate with brief periods of exposure to extreme heat, during which selection acted on capsid stability. The mutations that fixed in each line were identified, and the growth rate, decay rate, and fitness were measured at regular intervals throughout the adaptive walks.

Our first aim was to determine the extent to which pleiotropic effects on decay rate impede the long-term adaptation of growth rate, through a comparison of the adaptive trajectories of wildtype strains experiencing selection on both traits with the adaptation of the same strains first optimized under simple selection on growth rate and then experiencing complex selection on both traits. Additionally, we tested the hypothesis that complex organisms, or organisms experiencing complex selection acting on multiple traits, suffer reduced rates of adaptation, as predicted by evolutionary theory (Fisher 1930; Orr 2000; Welch and Waxman 2003). These aims have particular significance for understanding and predicting the adaptation of human and agricultural pathogens to novel hosts and environments as well as the persistence of preexisting adaptations in the face of novel selective conditions.

## Methods

### Experimental evolution

We performed complete experimental adaptive walks for two replicate lineages each of two pairs of wild-type and growth-adapted bacteriophage genotypes, ID8 and NC28, under selection on viral growth rate and capsid stability. All replicate lineages were initiated from unique sequence confirmed plaque isolates and evolved through serial flask passaging under a two-stage selection regime as described by McGee et al. (2016). For each passage, a culture of host cells (*Escherichia coli C*) was grown to a density of ~ 10^8^ cells/ml in 10 ml of lysogeny broth (10 g tryptone, 10 g NaCl, 5 g yeast extract per liter, supplemented with 2 mM CaCl) within a 125 ml Erlenmeyer flask at 37° in an orbital shaking water bath set to 200 rpm. The culture was then innoculated with ~ 10^5^ phage and allowed to propagate for 40 minutes, reaching a density of ~ 10^8^-10^10^ PFU/ml. Growth was then terminated by the addition of CHCl_3_ to lyse host cells, followed by centrifugation.

Heat shocks were performed as described by McGee et al. (2016). Briefly, aliquots of supernatant in microcentrifuge tubes were placed into an ice bath for five minutes to normalize temperatures, then placed in hot beads in a heating block set for 80° C and incubated for five minutes for ID8 lineages and four minutes for NC28 lineages, then transferred back to the ice bath for 5 minutes to terminate the heat shock. The duration times for the heat shock were calibrated for each genotype to generate the strongest possible reduction in population size, and thus the strongest possible selection on capsid stability, that the initial genotypes could handle without total population death. Population sizes were monitored by plating on agar plates prior to growth, immediately after growth, and immediately after heat shock. Population change rates were calculated on a log2 scale resulting in values of population doublings/halvings per hour. Adaptation was continued in this manner until no new mutations fixed in each population and average passage growth and decay rates remained stable for at least 20 passages.

ID8 and NC28 are ssDNA bacteriophages belonging to the family *Microviridae* and having genomes of 5540 nt and 6065 nt, respectively, and non-enveloped, tailless capsids with icosahedral geometry. Both phages were originally isolated from natural populations by Rokyta et al. (2006) and adapted to lab culture conditions on *E. coli C* by Rokyta et al. (2009). The growth-adapted strain of ID8 used in this experiment was the ID8a lineage described by Rokyta et al. (2009). Isolates confirmed to contain the full suite of published growth-rate mutations (Table 2) were used to initiate the ID8evol and NC28evol lineages in this experiment. The wild-type strains used to initiate the ID8wt and NC28wt lineages were taken from the same isolates described by Rokyta et al. (2006) and were sequenced to confirm they were free of additional mutations.

**Table 1.** List of mutations the fixed during the initial growth adaptation of ID8 and NC28.

| **Lineage** | **Prot. function** | **Prot. name** | **Aa position** | **ΔAa** | **Nuc. position** | **ΔNuc.** |
| --- | --- | --- | --- | --- | --- | --- |
| ID8 | DNA Packaging | C | 80 | N → S | 1958 | A → G |
| | Coat | F | 355 | P → S | 3628 | C → T |
| | Spike | G | 171 | T → A | 4493 | A → G |
| | DNA Pilot | H | 82 | T → A | 4770 | C → T |
| | DNA Pilot | H | 135 | H → Y | 4930 | G → A |
| NC28 | Scaffolding(Lysis) | D(E) | 129(69) | A(K) → S(N) | 2903 | G → T |
| | Coat | F | 272 | silent | 3904 | A → G |
| | DNA Pilot | H | 10 | V → A | 5116 | T → C |

A list of all mutations that fixed during the initial growth optimization of ID8 and NC28 performed by Rokyta et al. (2009), including the nucleotide and amino acid changes and a description of the affected proteins. Certain regions of the bacteriophage genome have overlapping genes, indicated with the second affected gene in parentheses.

**Table 2.**
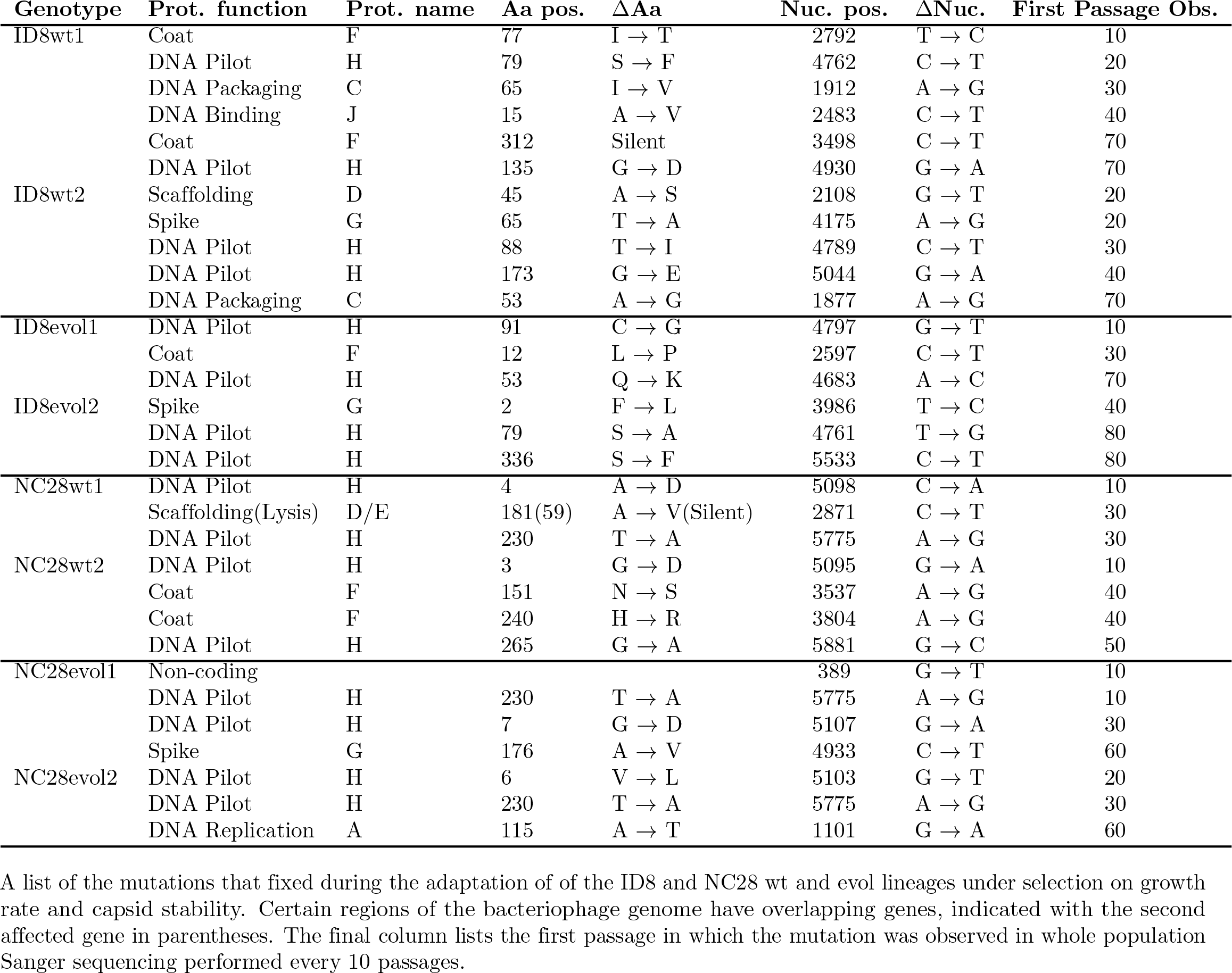
Mutations fixed during long-term adaptation of ID8 and NC28 lineages.

| Genotype | Prot. function | Prot. name | Aa pos. | ΔAa | Nuc. pos. | ΔNuc. | First Passage Obs. |
| --- | --- | --- | --- | --- | --- | --- | --- |
| ID8wt1 | Coat | F | 77 | I → T | 2792 | T → C | 10 |
| | DNA Pilot | H | 79 | S → F | 4762 | C → T | 20 |
| | DNA Packaging | C | 65 | I → V | 1912 | A → G | 30 |
| | DNA Binding | J | 15 | A → V | 2483 | C → T | 40 |
| | Coat | F | 312 | Silent | 3498 | C → T | 70 |
| | DNA Pilot | H | 135 | G → D | 4930 | G → A | 70 |
| ID8wt2 | Scaffolding | D | 45 | A → S | 2108 | G → T | 20 |
| | Spike | G | 65 | T → A | 4175 | A → G | 20 |
| | DNA Pilot | H | 88 | T → I | 4789 | C → T | 30 |
| | DNA Pilot | H | 173 | G → E | 5044 | G → A | 40 |
| | DNA Packaging | C | 53 | A → G | 1877 | A → G | 70 |
| ID8evol1 | DNA Pilot | H | 91 | C → G | 4797 | G → T | 10 |
| | Coat | F | 12 | L → P | 2597 | C → T | 30 |
| | DNA Pilot | H | 53 | Q → K | 4683 | A → C | 70 |
| ID8evol2 | Spike | G | 2 | F → L | 3986 | T → C | 40 |
| | DNA Pilot | H | 79 | S → A | 4761 | T → G | 80 |
| | DNA Pilot | H | 336 | S → F | 5533 | C → T | 80 |
| NC28wt1 | DNA Pilot | H | 4 | A → D | 5098 | C → A | 10 |
| | Scaffolding(Lysis) | D/E | 181(59) | A → V(Silent) | 2871 | C → T | 30 |
| | DNA Pilot | H | 230 | T → A | 5775 | A → G | 30 |
| NC28wt2 | DNA Pilot | H | 3 | G → D | 5095 | G → A | 10 |
| | Coat | F | 151 | N → S | 3537 | A → G | 40 |
| | Coat | F | 240 | H → R | 3804 | A → G | 40 |
| | DNA Pilot | H | 265 | G → A | 5881 | G → C | 50 |
| NC28evol1 | Non-coding | | | | 389 | G → T | 10 |
| | DNA Pilot | H | 230 | T → A | 5775 | A → G | 10 |
| | DNA Pilot | H | 7 | G → D | 5107 | G → A | 30 |
| | Spike | G | 176 | A → V | 4933 | C → T | 60 |
| NC28evol2 | DNA Pilot | H | 6 | V → L | 5103 | G → T | 20 |
| | DNA Pilot | H | 230 | T → A | 5775 | A → G | 30 |
| | DNA Replication | A | 115 | A → T | 1101 | G → A | 60 |

A list of the mutations that fixed during the adaptation of of the ID8 and NC28 wt and evol lineages under selection on growth rate and capsid stability. Certain regions of the bacteriophage genome have overlapping genes, indicated with the second affected gene in parentheses. The final column lists the first passage in which the mutation was observed in whole population Sanger sequencing performed every 10 passages.

### Sequencing and fitness assays

Each lineage was sequenced every 10 passages by amplifying the genome with two PCRs and sequencing the entire genome via Sanger sequencing. Whole population sequencing allowed detection of mutations that had fixed or reached high frequency. Each population-level sequence was compared to the sequence of its starting isolate and new mutations that fixed were recorded (Table 2).

At the end of adaptation, population fitness was assayed at several time points for each lineage over five replicates for intermediate time points and 8-10 replicates for the starting isolates and final time points of each population. For intermediate and final populations, a fresh population was grown for one passage starting from the −80° C freezer stocks of the end of growth passage of the time point before the desired assay time point. Aliquots of these fresh populations were used for assays, following the same growth period and heat-shock conditions described above.

Because our phage populations grow continuously rather than in discrete generations, we used Malthusian fitnesses (i.e., rates) rather than more typical Wrightian fitnesses (i.e., numbers of offspring). We measured growth rate, *γ*, during the 40 minute growth period, and decay rate, *δ*, during each heat-shock period (four minutes for NC28 and five minutes for ID8). Overall fitness, *ω*, was calculated as *ω* = *γτ*_*g*_ − *δτ*_*d*_, a combination of the growth and decay rates and the time spent during the growth period, *τ*_*g*_, and the time spent during heat shock *τ*_*d*_.

### Statistical analyses

Pairwise comparisons and sequential Bonferroni corrections were used to compare growth rate, decay rate, and fitnesses between time points and lineages. All statistical analyses were performed using R (R Development Core Team 2010).

### Data and reagent availability

Strains are available upon request. The GenBank accession numbers for ID8 and NC28 are DQ079898 and DQ079875, respectively, and all mutations are provided in Tables 1 and 2. The authors affirm that all data necessary for confirming the conclusions of the article are present within the article, figures, and tables.

## Results and Discussion

### Simultaneous improvement of putatively antagonistic phenotypes

We performed adaptive walks for two replicate lineages of wild-type and growth-adapted populations of microvirid phage strains ID8 and NC28, named ID8wt, ID8evol, NC28wt, and NC28evol, under conditions selecting on both phage growth rate under standard growth conditions (measured as population doublings per hour) and capsid stability (measured as viral decay rate, or population halvings per hour) under heat-shock con-ditions at 80° C (see methods). The initial ID8evol and NC28evol genotypes were the result of previous experimental growth-rate optimization performed by Rokyta et al. (2009) and were presumed to be at adaptive pleateaus with regard to growth rate prior to experiencing complex selection, and were selected for the experiment without any knowledge regarding their stability. Both lineages thus began with significantly higher growth rates than the ID8wt and NC28wt strains. ID8evol initially had a higher (worse) decay rate than the wild-type genotype, indicating that the initial growth adaptation of ID8 had negative pleiotropic effects on decay rate (Figure 1). Conversely, the NC28evol strain began with a significantly lower (better) decay rate than NC28wt, indicating that the original growth optimization of NC28 performed by Rokyta et al. (2009) had beneficially pleiotropic effects on decay rate.

**Figure 1.**
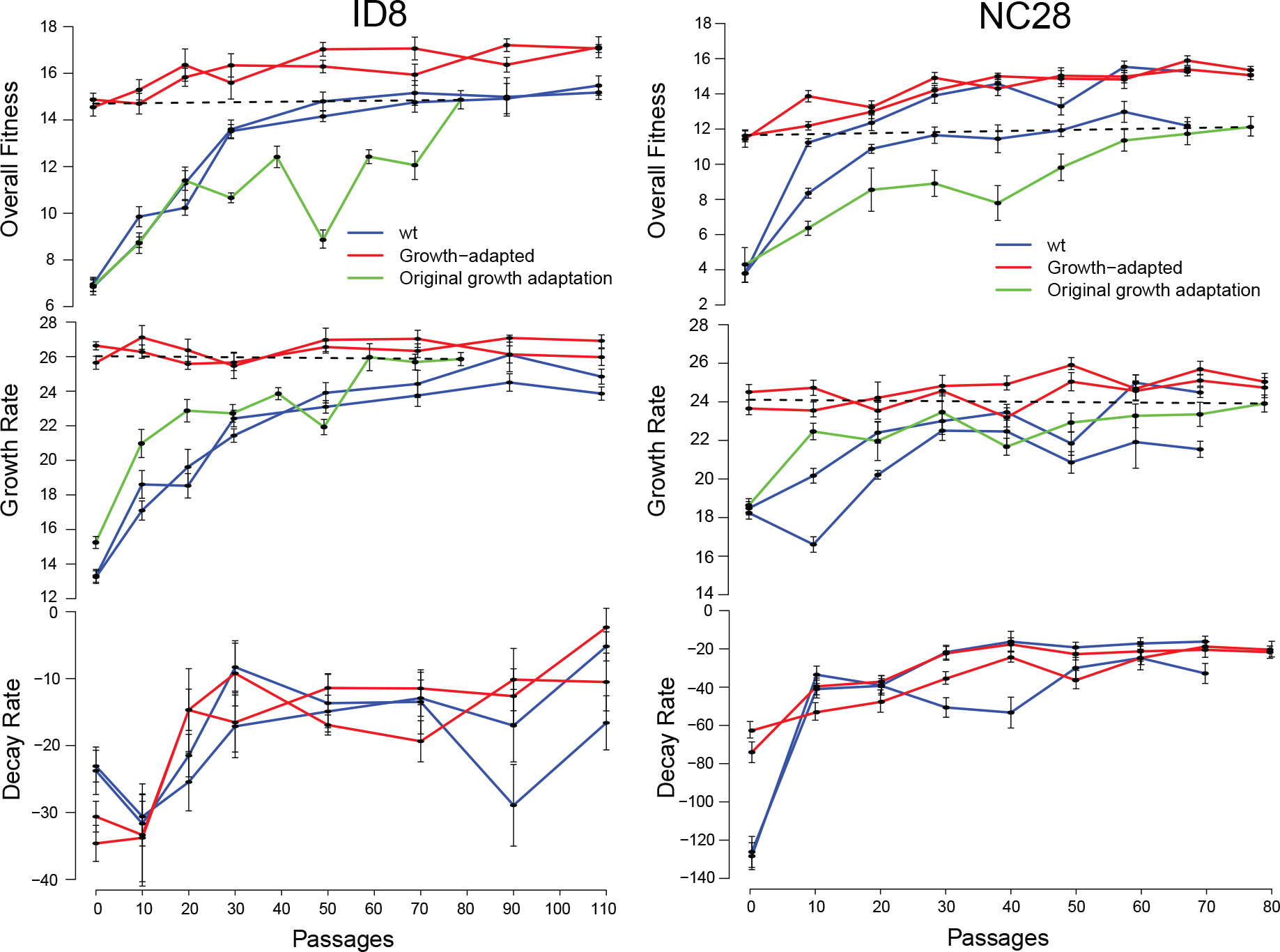
Fitness trajectories over an adaptive walk. Overall fitness (*ω*), growth-rate (*γ*), and decay-rate (*δ*) trajectories for ID8 and NC28 wild-type and growth-adapted lineages from the starting isolates (passage zero) to the end of adaptation. The overall fitness and growth-rate trajectories are included for the original growth adaptations of ID8 and NC28 (Rokyta et al. 2009), the endpoints of which were used to initiate the growth-adapted “evol” lineages. The fitness trajectories are significantly lower than those of the wild-type heat-shock trajectories, indicating that the original adaptations of ID8 and NC28 under growth-only selection fixed mutations that may not have been favored under two-trait selection.

**Figure 2.**
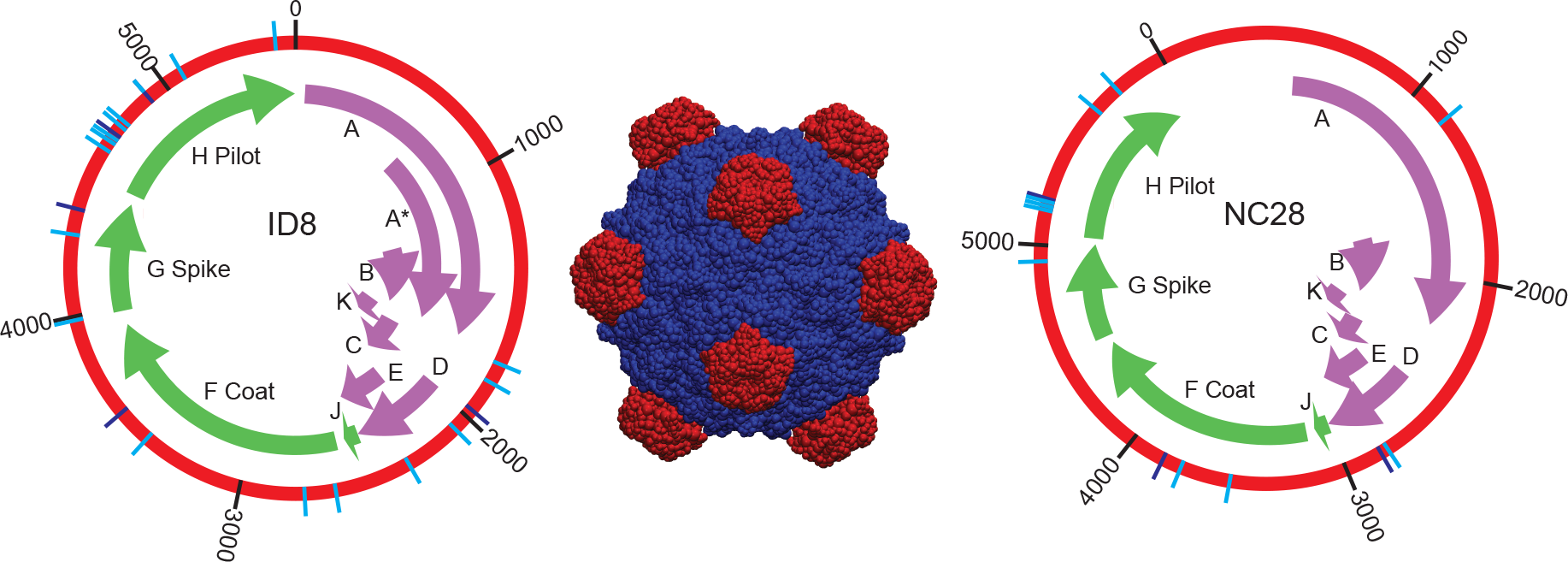
Map of fixed mutations. Left and right: Map of the locations of the mutations fixed during heat-shock adaptation of the wild-type (light blue) and growth-adapted (dark blue) lineages with each gene labeled. Structural genes present in the capsid are labeled in green. The gene with greatest activity in synergistic improvement of growth rate and stability is the gene encoding the DNA pilot protein H, with mutations also fixing in the DNA replication protein A, DNA packaging protein C, scaffold protein D, lysis protein E, capsid protein F, spike protein G, and DNA binding protein J. For additional details on each mutation, see Table 2. Center: A complete copy of the capsid with capsid protein F in blue and spike protein G in red.

Adaptation was continued for each lineage until they experienced a period of at least 20 passages with no improvement in either phenotype or fitness and with no new fixation of mutations, as determined by whole population Sanger sequencing every 10 passages. All four ID8 lineages (two replicates each of ID8wt and ID8evol) adapted for 110 passages, or ~330 generations. The NC28wt lineages adapted for 70 passages, and the NC28evol lineages adapted for 80 passages.

Both replicates of the ID8wt populations showed improvement of both growth rate and decay rate over the course of their adaptive walks, (Table 3; Figure 1). Remarkably, growth rate in the ID8evol lineages remained unchanged from the start to the end of adaptation despite significant improvements to decay rate, resulting in significantly improved overall fitness in both ID8evol replicates (Figure 1). Both NC28wt lineages similarly improved both traits over the course of adaptation. Growth rate improved significantly (though still by a small amount) in NC28evol2, and decay rate and overall fitness improved significantly in both NC28evol lines (Table 3; Figure 1). Mutations fixed in eight of 11 genes, with the vast majority of mutations affecting structural proteins F, G, and H (the coat, spike, and pilot proteins, respectively), and with a few exceptions growth rate and decay rate generally both improved throughout each complete bout of wild-type adaptation, indicating that most mutations were probably synergistically beneficial (Figures 1; 2).

**Table 3.** Initial and final growth rate, decay rate, and overall fitness for each lineage.

| **Lineage** | $\gamma_0$ | $\gamma_{evol}$ | $\delta_0$ | $\delta_{evol}$ | $\omega_0$ | $\omega_{evol}$ |
| --- | --- | --- | --- | --- | --- | --- |
| ID8wt1 | 13.32 ± 0.37 | 23.86 ± 0.38 | -23.08 ± 2.37 | -5.19 ± 2.17 | 6.96 ± 0.30 | 15.47 ± 0.41 |
| ID8wt2 | 13.24 ± 0.37 | 24.84 ± 0.43 | -23.77 ± 3.55 | -16.61 ± 4.04 | 6.85 ± 0.35 | 15.18 ± 0.30 |
| ID8evol1 | 25.65 ± 0.38 | 25.97 ± 0.40 | -30.62 ± 2.29 | -2.35 ± 2.85 | 14.55 ± 0.38 | 17.12 ± 0.45 |
| ID8evol2 | 26.63 ± 0.25 | 26.91 ± 0.38 | -34.62 ± 2.69 | -10.51 ± 4.31 | 14.87 ± 0.28 | 17.07 ± 0.17 |
| NC28wt1 | 18.48 ± 0.34 | 24.48 ± 0.26 | -127.74 ± 7.02 | -15.91 ± 2.84 | 3.80 ± 0.50 | 15.26 ± 0.24 |
| NC28wt2 | 18.24 ± 0.33 | 21.54 ± 0.42 | -125.50 ± 8.07 | -32.50 ± 5.22 | 3.80 ± 0.50 | 12.19 ± 0.47 |
| NC28evol1 | 24.50 ± 0.40 | 24.74 ± 0.52 | -73.58 ± 5.40 | -21.41 ± 3.19 | 11.43 ± 0.46 | 15.06 ± 0.26 |
| NC28evol2 | 23.65 ± 0.37 | 25.04 ± 0.37 | -62.27 ± 4.31 | -20.12 ± 4.10 | 11.62 ± 0.34 | 15.35 ± 0.21 |

The start and end measurements for growth rate, decay rate, and overall fitness are given for each lineage. Each value was averaged over 10 replicate fitness assays, and standard errors are given for each measurement.

We hypothesized that antagonistic pleiotropy would prevent the long-term simultaneous optimization of growth rate and capsid stability, based on the prediction that most mutations, particularly those involved with the improvement of function, are destabilizing (DePristo et al. 2005; Tokuriki et al. 2008; Tokuriki and Tawfik 2009; Phillips et al. 2017; Geller et al. 2018), especially when considering the effect of fixing multiple successive amino acid substitutions (Bloom et al. 2005; gong et al. 2013), and based on the prediction that optimal binding affinity for assembly may conflict with overall capsid stability (Zlotnick 1994; Ceres and Zlotnick 2002; Zlotnick 2003, 2005). Some empirical evidence has supported the hypothesized tradeoff between growth rate and stability. Dessau et al. (2012) identified a mutation in bacteriophage Φ6 that improved stability in a novel temperature environment but imposed a cost on viral reproduction. Heineman and Brown (2012) similarly found that mutations that improved survival of phage T7 in 6M urea came with large deleterious effects for growth rate. McGee et al. (2014, 2016) previously showed that when selection acts strongly on both traits, single-step mutations are available to wild-type ID8 that improve both traits simultaneously. However, the average benefit of each mutation to growth rate was significantly lower than when selection acted only on growth rate, and we did not know how the traits would interact over multiple mutational steps. It was therefore surprising that all wild-type lineages exhibited marked improvement in both components of fitness (with the exception of decay rate for ID8wt2, which did not change significantly in either direction), and that the ID8evol and NC8evol lineages improved decay rate so significantly with no deleterious pleiotropic effects on growth rate (in fact, growth rate improved slightly in NC28evol2).

Tradeoffs between traits have often emerged in organisms adapting to complex environments with multiple selection pressures, particularly when populations that have been locally adapted to a novel environment are compared to an ancestral population in an alternate environment (Travisano and Lenski 1996; Cooper and Lenski 2000; Bull et al. 2000; Cooper et al. 2001; Anderson et al. 2012; Oakley et al. 2014). This type of trade-off resulting from environmental complexity can be equated in our system to differences in selective complexity and the number of traits under selection. A similar result can be seen in the differential effects of the original growth-rate adaptation of ID8 and NC28, performed by Rokyta et al. (2009), on decay rate. Growth adaptation of ID8 resulted in a decay rate that was higher (worse) than the decay rate of wild-type ID8. However, growth adaptation of NC28 actually resulted in a decay rate that was half that of wild-type NC28 (Table 3; Figure 1). Selection during growth adaptation was blind to effects on decay rate, resulting in deleterious effects for decay rate during the original adaptation ID8, but not for NC28.

Our current experiment differs from others, because we alternated rapidly between selection on growth rate and stability, effectively selecting on both traits simultaneously. Bono et al. (2017) showed that tradeoffs from antagonistic pleiotropy emerge significantly more often in experiments with homogenous environments than in heterogenous environments, i.e., in instances where the traits incurring tradeoffs are not directly under selection during adaptation, and that tradeoffs were least likely in environments that were temporally heterogenous such that populations experienced frequent temporal shifts in the sets of traits under selection. Our results supported these findings, as long-term selection on growth rate alone without regard to effects on stability resulted in deleterious tradeoffs for stability in ID8, but populations experiencing simultaneous selection on both traits avoided deleterious effects on capsid stability by fixing mutations that improved both growth rate and decay rate. Experiments that fail to test for antagonism when selection acts simultaneously on all traits of interest may therefore identify antagonistic mutations simply because selection is not actively encouraging exploration of potential synergistic mutations, thereby lending unfounded support to the theoretical cost of complexity.

### Differential adaptive peaks arising from mutational contingency

At the end of adaptation, the growth rates and fitnesses of both ID8wt populations were significantly lower than the average growth rate of the ID8evol populations (two-sided Welch twosample *t*-tests, *p* < 0.01 for ID8wt1, *p* = 0.035 for ID8wt2, Bonferroni corrected for multiple comparisons, Table 3, Figure 1). The growth rate and fitness of NC28wt2 was also significantly lower than those of either NC28evol population (two-sided Welch two-sample *t*-test, *p* < 10^−3^ for all comparisons), but NC28wt1 achieved similar fitness and growth rate to both NC28evol lineages. Though significant, the disparities between the evol lineages and three of their counterpart wild-type lineages were small relative to the total fitness gains of the ID8wt and NC28wt lines.

In total, the ID8evol and NC28evol lineages experienced an additional 80 and 90 passages of adaptation, respectively, compared to the ID8wt and NC28wt lines, but given that the wt lines were continued until no new beneficial mutations were identified in Sanger sequences for at least 20 passages (40 passages for the ID8wt lines), we do not believe that these endpoint growthrate differences are attributable to the differences in passaging time. The end-point high growth rate achieved by NC28wt1 is further evidence that the disparity in time of adaptation is not responsible. Alternatively, at first glance the lower fitness peaks reached by three of the wild-type lineages may seem to be a consequence of antagonistic pleiotropy during evolution only under complex selection compared to evolution first under simple and then complex selection (as experienced by the evol lineages). However, the suboptimal peaks reached by the wild-type lines appear to be largely a consequence of historical contingency via the fixation of different mutational steps early in adaptation leading to alternate, higher adaptive peaks.

We found that the original growth adaptations of ID8 and NC28 involved mutational steps of significantly lower fitness effect (when assayed under two-trait selection, rather than the growth-only selection they experienced) than those taken by the ID8wt and NC28wt populations (Figure 1). These early mutations initiated trajectories which led to higher end-point growth rates than three of the four wt lines. Sign epistasis has been well documented within the the genomes of microvirid bacteriophages (Sackman and Rokyta 2018; Rokyta et al. 2011; Caudle et al. 2014; Doore and Fane 2015), indicating that the fitness landscape is at least moderately rugged, and that the higher adaptive peaks reached by the ID8evol lines may have been inaccessible without the mutations fixed during the original growth adaptation of ID8, which would not have been as highly favored as the mutations fixed by the ID8wt lines because of their reduced fitness benefit in the context of two-trait selection (as seen by the fitness trajectories of the original growth adaptations of ID8 and NC28 in green on Figure 1 being significantly below those of their wt counterparts during the first 80 passages of adaptation).

In particular, the mutation at position 171 of the spike protein G, which fixed in the original growth adaptation of ID8—as well as in an additional parallel line performed by *Rokyta et al.* (2009)—but not in either ID8wt line (Tables 1; 2), has the third largest effect on growth rate of any of the 24 unique first-step mutations previously identified on the wild-type ID8 background, and was also the most commonly fixed first-step mutation under growth-only selection (McGee et al. 2016). However, its deleterious effect on decay rate results in only a modest total benefit to fitness under heat-shock conditions, and it would therefore have been unlikely to fix early in the evolution of the ID8wt lineages when mutations of larger synergistic effect were available. Although we do not know the order of fixation of mutations in the original growth-adapted ID8 lineage, the large effect of the mutation at G171 makes it likely to have fixed in an early passage of growth-only selection. This mutation is also known to interact in a negatively epistatic manner with other large-effect growth-rate mutations, making it unlikely to be beneficial after other mutations fix on the ID8 wild-type background (Sackman and Rokyta 2018).

The fitness reached by the ID8evol lines was therefore likely only attainable by fixing mutations—including the mutation at G171—that would have been suboptimal under two-trait selection, meaning that, under this explanation, the elevated fitness of the ID8evol lines would be result of their sequential selection— experiencing first one-trait and then two-trait selection resulted in higher fitness than two-trait selection over the entire adaptive walk—rather than a result of antagonistic pleiotropic constraints acting on the ID8wt lines. Sequential selection allowed the exploration of regions of sequence space that were less accessible under simultaneous selection, as selection on a second trait altered the topology of the genotype-fitness landscape.

### No evidence for a cost of complexity

Rokyta et al. (2009) performed 80 passages of adaptation for the growth-adapted ID8 and NC28 lineages used in this experiment. Based on the stopping criteria used in their experiment and ours (passaging was stopped after no new mutations or fitness increases were detected for 20 passages), we can consider the ID8wt and NC28wt lineages completed at 90 and 80 passages, respectively (the ID8wt lineages were taken out to 110 out of an abundance of caution, but fitness plateaued at passage 70). The average rate of adaptation for the original growth-rate adaptations of ID8 and NC28, measured as the change in growth rate per ten passages, was 1.21 and 0.53 doublings per hour per ten passages, respectively, or 0.92 and 0.40 doublings per passage per ten passages, when scaled to the 45 minute total passage duration used to measure fitness in the heat-shock lineages. The average rate of growth-rate adaptation was higher (though not significantly) in the ID8wt and NC28wt lines (1.23 and 0.58 doublings per hour per ten passages, respectively), and their average rate of adaptation with regard to fitness was also higher (0.94 and 1.24 doublings per passage per ten passages, respectively, Figure 1, two-sided Welch two- sample t-test, *p* < 0.01 for NC28).

Looking more specifically at the explosive bursts of adaptation over the first 30 passages, the growth-selected lines averaged an improvement of 2.42 doublings per hour per ten passages, less (though not significantly so) than the rate of average growth improvement seen in the complex-selection lines, which was 2.51 doublings per hour per ten passages. By multiple measures of the rate of adaptation, the populations experiencing complex selection on average adapted at least as quickly—and in one case significantly more quickly—than those evolving in a simple environment.

Fisher’s geometric model (Fisher 1930) predicts that as the number of traits under selection increases, pleiotropy between traits increases the likelihood that a mutation of a given phenotypic magnitude is deleterious. A cost of complexity must therefore manifest as an inverse relationship between an organism’s rate of adaptation and the number of traits under selection (Orr 2000; Welch and Waxman 2003). These predictions are formulated in terms of organisms with differential numbers of phenotypic traits, but are equally relatable to organisms with the same number of phenotypic traits but with selection acting on a different subsets of those traits (McGee et al. 2016). Orr (2000) determined that the rate of adaptation should decline at a rate of 1/*n*, where *n* is the number of independent characters or traits under selection. We therefore predicted that adaptation under simultaneous selection on growth rate and capsid stability would be slower relative to growth-only adaptation, and that adaptation would be accomplished through mutations of smaller average effects.

We expected that much of long-term adaptation would result from compensation for deleterious pleiotropic effects, likely typified by sequential fixation of mutations of alternating signs of effect for each trait, with a smaller average benefit to total fitness per mutational step and thus a longer adaptive walk. Instead, however, adaptation required a very similar number of steps for optimization of two traits as were required for optimization of a single trait (growth rate). ID8wt2 fixed the same number of mutations as the original growth adaptation of ID8, and ID8wt1 only fixed one more mutation (one of which was silent and likely hitchhiked to high frequency; Tables 1; 2). NC28wt1 likewise fixed the same number of mutations as the original growth adaptation of NC28, and NC28wt2 fixed one additional mutation. We can be reasonably certain that, except in the case of hitchhiking, the mutations that fixed in each population are adaptive, given that the transit time of mutations of neutral or small effect should exceed the total time of the experiments, let alone the observed average fixation time of 30 − 60 generations (Kimura 1980).

These genotypes performed adaptive walks and achieved nearly the same (and in the case of NC28wt1, the same) high level of growth rate and produced (with the exception of ID8wt2) significant improvements in decay rate while taking almost the same average number of mutational steps as were required for optimization of growth rate alone. Additionally, the rate of growth-rate improvement per ten passages of adaptation was at least as high in the ID8wt and NC28wt populations as in their respective original growth-rate adaptations, despite experiencing selection on and improving a second trait. The rate of fitness improvement was slightly higher for ID8wt than during growth adaptation of ID8 and was three times as high in NC28wt as it was during NC28 growth adaptation. The higher rates of adaptation over adaptive walks of similar lengths imply that rather than suffering a reduced rate of adaptation resulting from antagonistic pleiotropy, synergistic pleiotropy facilitated a larger average fitness benefit per mutational step in the wild-type populations.

McGee et al. (2016) suggested that beneficial fitness effects that were observed for first-step mutations in wild-type ID8 under strong selection on growth and stability were larger than observed under growth-only selection or weak secondary selection on stability because populations experiencing strong selection began further from the adaptive optimum, thus erasing the cost of complexity. This followed from predictions from Fisher’s geometric model that average beneficial effects of mutations will be larger for populations farther from the optimum (Hartl et al. 1985). This argument does not hold, however, for our complete adaptive walks. Although the wild-type lineages started further from the optimum under complex selection than under growth selection, this difference in selection strength would have been erased as both sets of populations approached their respective optima. The wild-type populations did not exhibit any cost of complexity, as they began adaptation further from the optimum than they did under growth adaptation, but reached a fitness plateau over a similar number of mutational steps and similar number of generations with a higher rate of adaptation. Wagner et al. (2008) showed that quantitative trait loci (QTL) in mice on average only affect a small subset of all traits, and that mutational effect sizes did not decrease with an increasing level of pleiotropy, suggesting that modularity of pleiotropic interactions can alleviate the cost of complexity. Our results extended this finding of a nonexistent cost of complexity even to cases where mutations do affect all traits under selection simultaneously, as is the case for mutations affecting assembly kinetics and stability of structural capsid proteins.

## Conclusions

Theoretical and empirical results demonstrate that the majority of mutations improving protein function are energetically destabilizing to some extent, resulting in tradeoffs between stability and function. In viruses, mutations that increase the binding affinities between capsid subunits (i.e., stability) are predicted to disrupt capsid assembly and thus impose costs on viral growth rates. Contrary to our expectations, we found that the optimization of growth rate was not hindered by pleiotropic effects on stability. All wild-type heat-shock lineages achieved large gains in growth rate, while capsid stability was also improved significantly in all but one population, with no deleterious pleiotropic effects on growth rate observed in lines that were already optimized for growth rate. Remarkably, adaptation to a complex selective environment on average required less than one additional mutation relative to adaptation in a simple environment, the time required for adaptive walks was not longer, and the rate of fitness increase per unit of time was at least as high, and sometimes higher, in a complex environment. These results conflict with the theoretical prediction that increasing selective or organismal complexity is inversely related to the rate of adaptation.

We concluded that our experimental lineages were able to avoid tradeoffs and a subsequent reduction in the rate of adaptation by fixing mutations that were synergistically pleiotropic, and the diversity of substitutions fixed by each replicate lineage indicates that a large number of this class of mutation are available to adapting populations. Additionally, the properties of adaptive walks were strikingly similar across genotypes and selective environments, in terms of the number of mutations fixed, the number of generations required, and the overall rate of adaptation. These results imply that pathogenic viruses adapting to new hosts and highly variable environments may be able to adapt far more quickly than expected by fixing beneficial mutations of large effect that simultaneously improve multiple traits.

## Acknowledgements

Funding for this work was provided by the U.S. National Institutes of Health (NIH) to D.R.R. (NIH R01 GM099723).

